# Nuclear genetic diversity of head lice sheds light on human dispersal around the world

**DOI:** 10.1101/2022.08.09.503408

**Authors:** Marina S. Ascunce, Ariel C. Toloza, Angélica González-Oliver, David L. Reed

## Abstract

The human louse, *Pediculus humanus*, is an obligate blood-sucking ectoparasite that has coevolved with humans for millennia. Given the intimate relationship between this parasite and the human host, the study of human lice has the potential to shed light on aspects of human evolution that are difficult to interpret using other biological evidence. In this study, we analyzed the genetic variation in 274 human lice from 25 geographic sites around the world by using nuclear microsatellite loci and female-inherited mitochondrial DNA sequences. Nuclear genetic diversity analysis revealed the presence of two distinct genetic clusters I and II, which are subdivided into subclusters: Ia-Ib and IIa-IIb, respectively. Among these samples, we observed the presence of the two most common louse mitochondrial haplogroups: A and B that were found in both nuclear Clusters I and II. Evidence of nuclear admixture was uncommon (12%) and was predominate in the New World potentially mirroring the history of colonization in the Americas. These findings were supported by novel DIYABC simulations that were built using both host and parasite data to define parameters and models suggesting that admixture between cI and cII was very recent. This pattern could also be the result of a reproductive barrier between these two nuclear genetic clusters. In addition to providing new evolutionary knowledge about this human parasite, our study could guide the development of new analyses in other host-parasite systems.

## Introduction

The human louse (*Pediculus humanus*, Phthiraptera: Anoplura) is a blood-sucking, wingless, human-specific ectoparasite that is obligate (cannot live without the host) and permanent (complete their life cycle on a host) [1]. This relationship has made humans and lice ideal for coevolutionary studies [2,3]. In the past, studies have shown that primate lice cospeciated with their hosts over the last 25 million years (MY) [2,3]. Furthermore, cophylogenetic analyses revealed that human and chimpanzee lice in the genus *Pediculus* diverged ca. 5.6 MY ago (MYA), which is contemporaneous with the known divergence of chimps and humans [4]. Human head lice have been found in archeological sites, including a head louse nit obtained from human remains in northeastern Brazil dated 10,000 years old [5] and a nit from Israel dated 9,000 years old [6] (Fig 1), among others. Multiple archeological findings of human lice around the world are shown in Fig 1 supporting the idea that lice are one of the oldest known human ectoparasites. In host-parasite systems, these long-term associations could lead to reciprocal adaptations in the hosts and parasites (i.e., coevolution), which allows one to build phylogenies for both the parasite and host and to obtain time estimations of events that co-occurred in both the host and parasite [7]. In cases where the parasite and host have an inextricable relationship, the parasite phylogeny could reflect host evolutionary history [8–10]. In some cases, there are also parasite-specific life history traits such as relatively faster evolutionary rates and larger population sizes that allow parasites to maintain greater levels of genetic diversity over time than their hosts. This allows researchers to harness the parasites’ genetic diversity in order to explore unclear aspects of host evolution based on direct host evidence, such as archeological remains or host molecular data [10–13]. Evolutionary studies of human lice have already proven themselves as a useful tool in human evolutionary studies. They have provided insights on the ecology of early humans, such as when humans started using clothes [14,15] and how humans travelled globally [16–18]. Recently, human DNA was extracted from nits found on hairs from ancient mummies in South America revealing ancestral human migration routes [19]. Moreover, almost 20 years ago, Reed et al. [2] described highly divergent human louse mitochondrial clades hypothesizing that each louse mitochondrial lineage evolved on different hominin hosts. The authors suggested that as anatomically modern humans (AMH) migrated out of Africa and interacted with other hominins, direct physical contact among hominins allowed different lineages of lice to be transferred from host to host [2]. The hypothesis of direct contact between modern and archaic hominins predicted using human louse data was later confirmed through host genomic studies comparing ancient DNA genomes from hominin remains with AMH genomes [20].

**Fig 1.** Humans and lice. The map shows the geographic distribution of the modern human head lice included in this study using green dots. Archeological findings of human lice are shown with the figure of a human louse on the map with the corresponding estimated dates from: [3,5,6,21,22]. In addition, the map reflects the approximate locations of hominin fossil remains and their proposed distribution based on: [23–38]. Each hominin is color coded as follows: Neanderthal (Blue), Denisovan (Black), and Anatomical Modern Humans (Orange). Map author: Maulucioni (https://commons.wikimedia.org/wiki/File:World_map_with_the_Americas_on_the_right.png). This file is licensed under the Creative Commons Attribution-Share Alike 4.0 International license. https://creativecommons.org/licenses/by-sa/4.0/legalcode.

The genetic diversity of human lice has been widely studied based on mitochondrial DNA. However, little is known about the nuclear genetic diversity of this human parasite, particularly on a global scale. Early human louse mitochondrial studies revealed the presence of three deeply divergent mtDNA clades or haplogroups named A, B and C [2,3,14,17,39]. Haplogroup A is the most common of the human louse haplogroups, has a worldwide distribution and has been found in lice from human archeological remains in Israel from 2,000 years ago [21]. Haplogroup B has been found in the Americas, Europe, Australia, North and South Africa [17,39–41], as well in ancient human remains in Israel [21] and Peru [3]. Haplogroup C has a more geographically restricted distribution and is only present in lice found in Nepal, Ethiopia, the Democratic Republic of the Congo, Pakistan and Thailand [14,21,42]. Recent studies have suggested the presence of three more clades: D, E and F [41,43,44]. Clade D is sister to clade A found in Ethiopia, Zimbabwe and the Democratic Republic of the Congo [41,43,45]. Clade E is sister to clade C and was detected in head lice from Senegal and Mali [41]. Clade F appears to be specific to South America, as it has not been reported elsewhere [44]. Head lice can be found in all the mitochondrial clades (A-F), whereas clothing lice are only found in clades A and D [45]. A recent study using all available cytochrome *b* (cyt*b*) gene human louse sequences corroborated the presence of these six clades: A, B, C, D, E, and F [46]. Their phylogenetic analysis indicated a first split around 1 MYA between clade ABDF and clade CE, additionally, clades AD and BF had the most recent common ancestor around 0.9 MYA, with later splits of clade D from A, F from B and E from C dating around the same period at 0.6 MYA [46]. Despite the extensive work that has been done by many researchers on louse mitochondrial diversity, there are some qualities that this maternally transmitted molecular marker cannot resolve such as hybridization events among different louse populations. To overcome this limitation 15 new nuclear microsatellite markers that have biparental inheritance have been developed [16]. In this new study we used the 15 microsatellite markers in order to provide a comprehensive understanding of human louse genetic diversity and to assess admixture among human lice.

In our previous study of louse microsatellite diversity, we only included 75 head lice from 11 geographic sites from four geographic regions [16]. We found that nuclear genetic diversity in human lice revealed continental differences and that the genetic pattern found in the Americas could be mirroring the human host colonization of the New World. In this new study, we expanded the sampling of lice to 274 and included 25 locations distributed over 10 geographic regions (Table 1, S1 Table, Fig 1). This broader sampling allows us to improve our understanding of human-louse nuclear genetic diversity while also assessing its correlation with mitochondrial diversity at a more global scale. This new study also addresses some of the questions uncovered in another work where we analyzed 450 mtDNA COX1 sequences from lice collected in the Americas [17]. Besides enlarging the sampling size and widening the geographic range in this study, we tested seven different evolutionary models by conducting DIYABC-simulations. For these new simulations, the parameters were developed with both host and parasite information. We wanted to explore the origin of the hybridization between these louse nuclear genetic clusters and test if the hybrid lice could be the result of the retention of ancient polymorphisms due to incomplete lineage sorting (ILS) (Model 1 and 2) or admixture among louse genetic clusters. For Models 1 and 2, we tested the retention of ancient polymorphisms due to incomplete lineage sorting (ILS) on the hybrid lice, where the hybrid lice were derived from lice that parasitized the heads of Neanderthals (Model 1), or the heads of anatomical modern humans (AMH) (Model 2). There were four scenarios considering different times for the admixture. An ancient admixture model tested hybridization between lice parasitizing Neanderthals and AMH during periods of contemporaneity of the two hosts (Model 3). Admixture during European colonization of the Americas 500 years ago was analyzed in Model 4. In Model 5, we tested the hybridization occurring in the last 100 to 40 years around World Wars I and II. Model 6 reflected the beginning of globalization from the 1980’s to ten years ago, while Model 7 referred to the last 10 years. In addition to providing new evolutionary knowledge about this parasite, our study could guide the development of new analyses in other host-parasite systems using this novel approach for the DIYABC-simulations.

**Table 1.** Microsatellite polymorphisms among human louse populations. Groups reflect their geographic region and STRUCTURE clustering: Africa-Israel-Spain (AIS), South and Southeast Asia (SAs), East Asia (EAs), Europe (E), North America (NA), Mesoamerica (MA), Central America (CA), and South America (SAm).

| Geographic Region | Group | Country | Site | N | $N_a$ | $H_O$ | $H_E$ | $F_{IS}$ | $P$-value |
| --- | --- | --- | --- | --- | --- | --- | --- | --- | --- |
| Africa | AIS | Rwanda | Rw | 1 | 1.533 | n/a | n/a | n/a | - |
| West Asia | AIS | Israel | Isr | 2 | 1.667 | n/a | n/a | n/a | - |
| Central Asia | SAs | Nepal | Np | 1 | 1.667 | n/a | n/a | n/a | - |
| South and Southeast Asia | SAs | Thailand | Th | 1 | 1.333 | n/a | n/a | n/a | - |
|  | SAs | Cambodia | Cam | 22 | 6.8 | 0.483 | 0.703 | 0.319 | 0.000 |
|  | SAs | Laos | Ls | 4 | 3.2 | n/a | n/a | n/a | - |
| East Asia | EAs | Mongolia | Mn | 10 | 5 | 0.524 | 0.706 | 0.267 | 0.005 |
| Europe | E | Turkey | Tur | 12 | 3 | 0.232 | 0.408 | 0.407 | 0.000 |
|  | E | Croatia | Croa | 6 | 1.133 | n/a | n/a | n/a | - |
|  | E | Norway | Nw | 1 | 1.067 | n/a | n/a | n/a | - |
|  | E | Netherlands | Neth | 64 | 2.267 | 0.032 | 0.248 | 0.863 | 0.000 |
|  | E | France | Fr | 3 | 1.133 | n/a | n/a | n/a | - |
|  | E | United Kingdom | UK | 27 | 2.533 | 0.145 | 0.329 | 0.550 | 0.000 |
|  | AIS | Spain | Spain | 3 | 2.067 | n/a | n/a | n/a | - |
| North America | NA | USA | Oce | 25 | 2.8 | 0.083 | 0.352 | 0.793 | 0.000 |
|  | NA | USA | SF | 1 | 1 | n/a | n/a | n/a | - |
|  | NA | USA | Martin | 1 | 1 | n/a | n/a | n/a | - |
|  | NA | USA | WPB | 1 | 1.467 | n/a | n/a | n/a | - |
|  | NA | USA | Nash | 18 | 2.467 | 0.037 | 0.311 | 0.864 | 0.000 |
|  | NA | USA | MC | 22 | 3.867 | 0.173 | 0.452 | 0.587 | 0.000 |
| Meso America | MA | Mexico | MX-1 | 18 | 3.13 | 0.208 | 0.396 | 0.481 | 0.000 |
|  | MA | Mexico | DF | 1 | 1 | n/a | n/a | n/a | - |
| Central America | CA | Honduras | Hon | 20 | 4.2 | 0.365 | 0.567 | 0.312 | 0.000 |
| South America | SAm | Peru | Peru | 1 | 1.267 | n/a | n/a | n/a | - |
|  | SAm | Argentina | Arg | 9 | 3.2 | 0.349 | 0.482 | 0.231 | 0.012 |
| Total/Average |  |  | 25 | 274 | 12 | 0.257 | 0.402 | 0.469 | 0.000 |

$N$ = sampling size; $N_a$ = Average number of alleles; $H_O$ = observed heterozygosity; $H_E$ = expected heterozygosity; $F_{IS}$ = index of inbreeding of individuals resulting from the non-random union of gametes within a subpopulation, positive value indicates heterozygote deficit; $P$ proportion of permutations leading to a $F_{IS}$ value larger or equal to the observed one. n/a indicates values not estimated due to sample size smaller than 9 individuals.

## MATERIALS AND METHODS

### Ethics statement

The Institutional Review Board of the University of Florida exempted the study from review (Exemption of Protocol #2009-U-0422) and waived the need for written informed consent of the participants. This exemption is issued based on the United States Department of Health and Human Services (HHS) regulations. Specifically, HHS regulation 45 CFR part 46 applies to research activities involving human subjects. Because louse removal was voluntary and no information was recorded that would allow patients to be identified directly or through identifiers linked to them, the University of Florida waived the need for written informed consent from the participants. Additional information about collection procedures has been included in the Supporting Information section.

### Sampling

Sampling details are provided in Fig 1, Table 1 and S1 Table. Briefly, 274 human head lice were collected from different individuals at 25 locations throughout the world covering 10 major geographic regions. A detailed protocol for collecting human lice has been included in the Supporting Information section.

### DNA extraction

We followed our DNA extraction protocol as detailed in [16,17] in which each individual louse was cut in half, one half was kept in 95% Ethanol for storage, while the other half was placed into 1.5 ml Eppendorf tubes containing a cell lysis and proteinase K solution and manually ground using a pestle for DNA extraction. DNA was extracted using the Puregene Core Kit A (QIAGEN, Valencia, California) following manufacturer protocols for tissues however only a third of the volume suggested in the protocol was used due to the small size of the louse material. DNA was quantified using a NanoDrop 1000 spectrophotometer (Thermo Scientific), and DNA samples were diluted to reach a concentration between 5 to 10 ng/μL.

### Multiple co-amplification: Multiplexes

A total of 15 microsatellite loci were amplified using four multiplexes as described in [16]. We used fluorescently labeled universal primers: M13 (5’-CACGACGTTGTAAAACGAC-3’) and CAG (5’-CAGTCGGGCGTCATCA-3’), each labeled with a unique fluorescent tag (e.g., FAM, VIC, NED, PET; Applied Biosystems) to co-amplify multiple loci. Locus-specific primers were modified by adding the matching 5′ universal primer sequence tails. The complete set of 15 loci were amplified in four separate 15 μL multiplex-PCR reactions containing 7.5 μL of 2X Master Mix (Type-It Microsatellite PCR kit, Qiagen, Venlo, Netherlands), 0.015-0.15 μL of unlabeled forward primer with tail (10 μM), 0.15-0.8 of unlabeled reverse primer (10 μM), 0.5 μL labelled universal tail primer (10 μM), 1-2 μL of total genomic DNA (10-20 ng) and dH_2_O added up to the final volume of 15 μL. Two thermal cycling profiles were used, both beginning with initial denaturation at 95°C (5 min) and ending with a final extension of 72°C (40 min). The v1 (touch-down PCR) protocol consisted of 10 cycles of 94°C (30 sec), 60°C/55°C (-0.5°C/cycle) for 45 sec, and 72°C (45 sec) followed by 25 cycles of 94°C (30 sec), 55°C (45 sec), and 72°C (1 min). The v2 protocol consisted of 35 cycles of 94°C (30 sec), 52°C (45 sec), and 72°C (45 sec). PCR products were electrophoresed on 1.5% and 4% agarose gels stained with ethidium bromide and visualized under ultraviolet light. Dilutions in dH_2_O from amplicons (1:100) were run on an ABI 3730xl 96-capillary sequencer using GeneScan 600 LIZ as an internal size standard (Applied Biosystems) at the University of Florida biotechnology facility (ICBR). Microsatellite genotypes were scored using the GeneMarker version 1.60 software (SoftGenetics, LLC).

### Mitochondrial PCR amplification and sequencing

The mitochondrial gene cytochrome *c* oxidase 1 (COX1) was amplified through polymerase chain reaction (PCR) using the primers H7005 and L6625 [9] as described in [2]. Each PCR reaction consisted of 25 μL total volume, which included 10 μL of MasterMix (5 PRIME), 1 μL of each primer, 2-4 μL of total genomic DNA, and water. The PCR’s cycling profile began with an initial denaturation at 94°C (10 min) followed by 10 cycles of 94°C (1 min), 48°C (1 min), and 65°C (2 min) (decreased by 0.5°C per cycle). Then, there were 35 cycles of 94°C (1 min), 52°C (1 min), and 65°C (2 min) and a final extension of 65°C (10 min). PCR products were run in agarose gel to verify amplification then purified using ExoSAP-IT (USB Corporation, Cleveland, Ohio). Purified PCR products were submitted for sequencing at the University of Florida DNA Sequencing Core Laboratory (ICBR, Gainesville, Florida) using standard fluorescent cycle-sequencing PCR reactions (ABI Prism Big Dye terminator chemistry, Applied Biosystems).

### COX1 sequence editing, alignment, and haplotype reconstruction

For each louse sample, the forward and reverse sequences were edited manually and aligned using Sequencher 4.5 (Gene Codes Corporation, Ann Arbor, Michigan). Consensus sequences were generated for each sample using both forward and reverse sequences. These new louse COX1 sequences were aligned along with previous sequences of lice used in our previous work [17]. A final alignment of 379 bp was used to construct a matrix of pairwise differences using uncorrected *p*-distances (proportion of nucleotide sites at which two sequences being compared are different) with MEGA 4 [47]. The genetic relationships were estimated by constructing neighbor-joining (NJ) trees [48] using PAUP* 4.0b10 [49] following Light et. al. [1]. Bootstrapping was performed using 100 pseudo-replications of the data set.

### Analyses of genetic diversity and population structure among collection sites using microsatellites

We used the software Arlequin 3.5.1.2 [50] to estimate population genetic diversity values based on the microsatellite data. For all sites regardless of the number of lice per site, we determined the number of alleles per locus. Then, for the 11 sites with sample sizes exceeding 8 lice per site, we also estimated observed and expected heterozygosity (H_O_ and H_E_), and the mean *F*_IS_ over loci per population with significance tested using 1,023 random permutations. For those 11 sites with sample sizes exceeding 8 lice per site, population pairwise *F_ST_* values were estimated using Arlequin 3.5.22 [51]. Significance values for pairwise *F_ST_* values were estimated using the permutation procedure with 1,000 permutations.

To understand how the human louse genetic diversity is distributed around the globe, we used three different methods that are based on the clustering of similar genotypes. First, we used STRUCTURE [52], a model-based Bayesian clustering method, that assumes Hardy–Weinberg and linkage equilibrium among loci. In addition, we used two multivariate methods, which do not carry these assumptions and could visualize potential cryptic structure, PCoA (Principal Coordinates Analysis) in GenAlEx 6.5 [53,54] and Discriminant Analysis of Principal Components (DAPC) [55].

#### Bayesian clustering analyses

The Bayesian clustering approach implemented in STRUCTURE 2.3.4 [52] uses the individual multi-locus genotypic data to evaluate models assuming different numbers of genetic clusters (*K*) based on the posterior probabilities given the data and model. Based on the individual multilocus genotypes and the allele frequencies estimated for the reconstructed clusters, each individual’s genome was probabilistically partitioned into membership fractions (*q*) (ancestry) in each cluster. Admixed ancestry is modeled by assuming that an individual *i* has inherited some fraction (*q*) of its genome from ancestors in population *k* [52]. Hybrids can then be inferred when observing an individual being intermediate of some degree between two or more clusters in terms of the *q*-value. For example, individuals with *q* = 0.5 would likely be first-generation hybrids. We identify possible hybrids based on a *q*-value threshold of 0.2 as suggested in VäHä and Primmer (2005) [56], where a threshold *q*-value of 0.2 means that individuals with *q*-value between 0 and < 0.20 or > 0.8 and 1 are classified as non-admixed and any individual with *q*-value between 0.2 and 0.8 are classified as hybrids. All simulations used 50,000 Markov chain Monte Carlo (MCMC) generations in the burn-in phase and 100,000 generations in the data collection phase. Ten independent runs were performed using default parameters for each *K*, to ensure equilibrium during burn-in and consistency in estimation of the posterior probabilities. Selection of the number of distinct clusters was based on the evaluation of the Δ*K* statistic [57] using the online tool STRUCTURE HARVESTER [58]. The ten STRUCTURE runs at each *K* produced nearly identical individual membership coefficients. The run with the highest likelihood of the data given the parameter values for the predominant clustering pattern (i.e. the mode) at each *K* was used for plotting with DISTRUCT [59].

#### Multivariate analysis using Principal Coordinates Analysis (PCoA)

This multivariate analysis was conducted using pairwise genetic differences between individuals calculated in GenAlEx 6.5 [53,54] to validate and further define genetic clusters found using STRUCTURE.

#### Multivariate analysis using Discriminant Analysis of Principal Components (DAPC)

This analysis was conducted using the ‘*adegenet’* package 2.1.3 [60], in the R software environment version 3.6.3 [61]. For these analyses we only included 175 lice from which we were able to obtain both microsatellite genotypes and mitochondrial haplogroup sequences. An additional DAPC included the same louse samples, however we considered four groups depending on nuclear clusters and haplogroup membership.

#### Allelic richness within each main nuclear STRUCTURE genetic cluster

We estimated the allelic richness within each main nuclear STRUCTURE genetic cluster using the rarefaction method implemented in the computer program ADZE (Allelic Diversity Analyzer) version 1.0 [62]. Because allelic richness measurements considered sample size, the rarefaction method allows the estimation of allelic richness for different random subsamples of size “g” from the populations [63,64].

#### Analyses based on STRUCTURE genetic clusters and mitochondrial clades

To gain a better understanding of the geographic distribution of the nuclear and mitochondrial diversity, we conducted a set of Kruskal–Wallis nonparametric analyses of variance tests [65]. We included the same 175 lice from which we were able to obtain both microsatellite genotype and mitochondrial haplogroup sequence data and the same eight geographic regions as described above. Calculated medians were compared and separated at *P* < 0.05 [66]. We compared the Observed vs. Expected Frequency *X*^2^ test to evaluate the fit of the data (frequencies) to any arbitrary set of expected frequencies among the eight geographic regions. All statistical tests were performed with a significance level of α=0.05.

Analysis of molecular variance (AMOVA) considering the number of different microsatellite alleles (*F*_ST_ based) was used to investigate the partitioning of genetic variation between the two nuclear clusters: cI and cII using Arlequin 3.5 [50]. The statistical significance of differentiation at each level was assessed by means of 1,023 permutations.

The historical migration patterns among the genetic clusters identified using the methods describe above were estimated in Migrate-n 3.3.0 using the maximum likelihood approach [67,68]. Migrate employed estimates of genealogies to sample areas of the coalescence space with the highest likelihood. The random number of seed and starting values of □ and 4N_e_m (where N_e_m is the indirect measure of gene flow) were based on *F*_ST_ as starting parameters and we used a Brownian mutation model. The data was analyzed using 10 short (10^4^ MCMC steps) and 3 long chains (10^5^ MCMC steps), together with adaptive heating to assure convergence.

#### Comparison of demographic models

We further inferred the demographic history of human lice, based on microsatellite data, using the approximate Bayesian computation (ABC) approach [69], implemented in the program DIYABC, v2.0 [70]. We tested different evolutionary Models in which each louse genetic cluster evolves on different human host populations and diverges in tandem with their hosts. A total of seven different evolutionary Models were considered with estimations of population size and divergence times based on information of both human evolution and lice (Table 2, Fig 2). The first three Models are directly testing Reed et al. (2004) [2] hypothesis of ancient hybridization between modern and archaic humans. Models 1 and 2 aimed to unravel the retention of ancient polymorphisms due to incomplete lineage sorting (ILS) on the hybrid lice, where the hybrid lice were derived from lice that parasitized the heads of Neanderthals (Model 1), or the heads of anatomical modern humans (AMH) (Model 2). In Model 3, admixture events between lice parasitizing Neanderthals and AMH during periods of host coexistence were tested. The last four models (Model 4, 5, 6 and 7) do not require ancient hominins, and are based solely on hybridization events within AMH during recent time periods. In Model 4, we tested the hypothesis that the hybrid lice were the product of hybridization between lice from Native Americans and the first Europeans during the European colonization of the Americas 500 years ago. In Model 5, we tested the hybridization occurring in the last 100 to 40 years around World Wars I and II. Model 6 reflects the beginning of globalization in the 1980’s to ten years ago, while Model 7 refers to very recent events in the last 10 years.

**Fig 2.** Evolutionary Models. We used seven different demographic historical models with three different level of parasitism and three different generational times for the lice that lead to a total of 63 different scenarios that were compared using the approach from the DIY-ABC program.

**Table 2.**
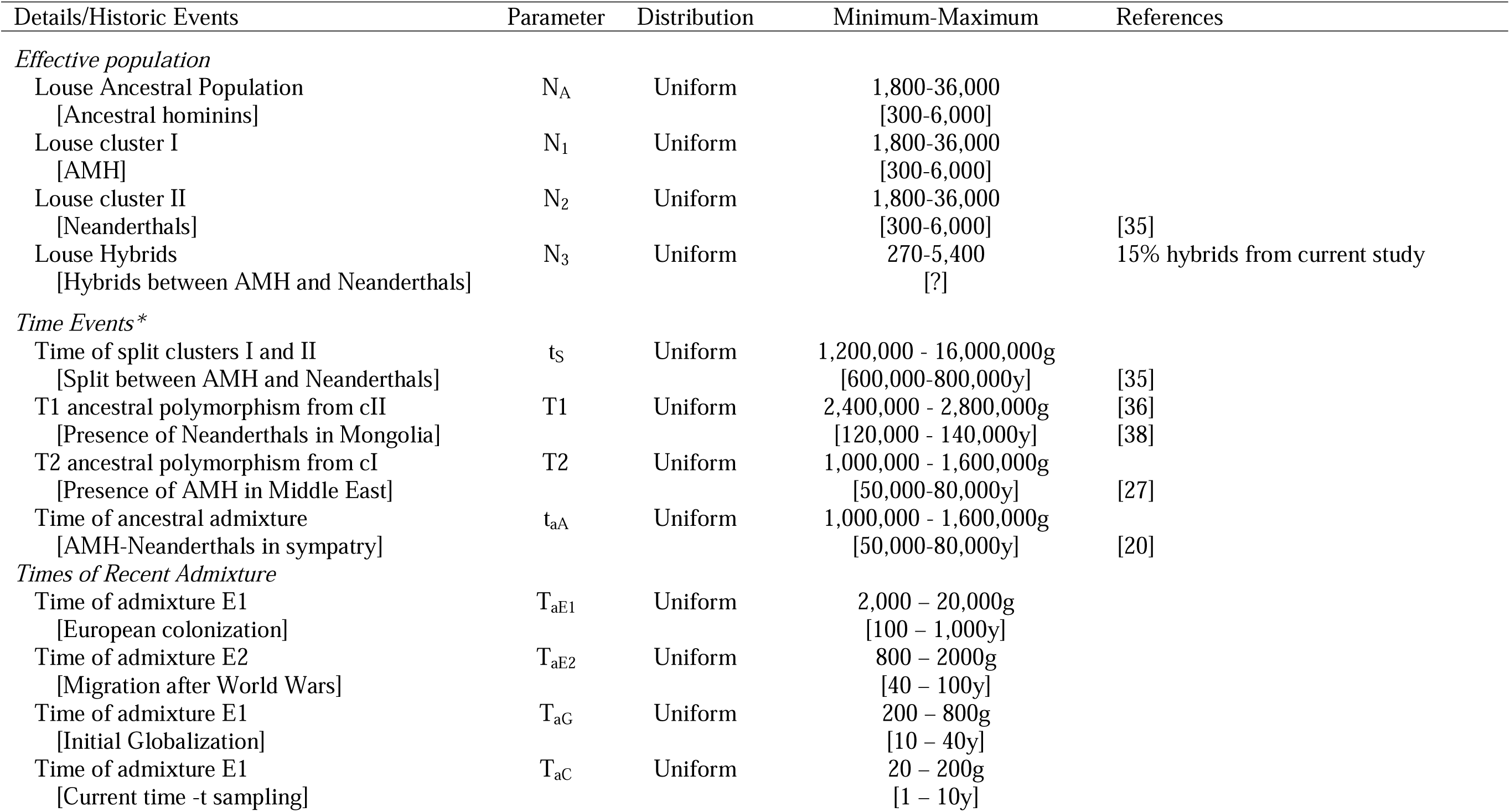
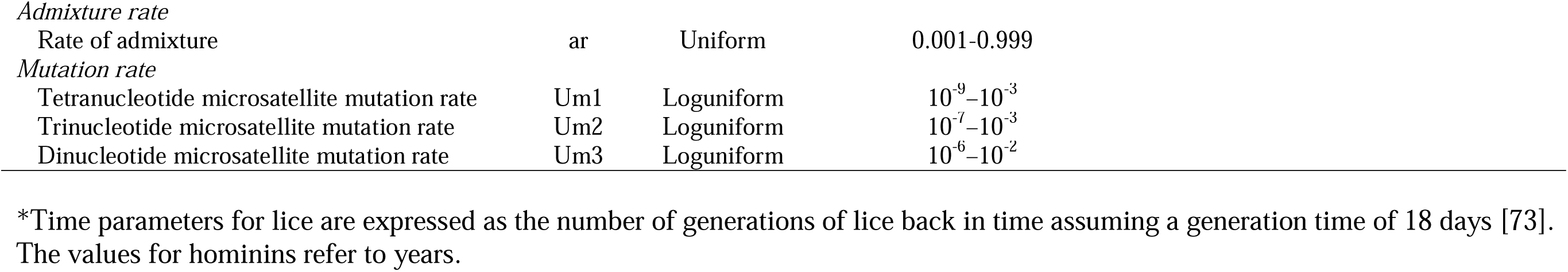
Prior distributions of historical and demographic parameters for the demographic scenarios. Between the brackets [] are the host values obtained from anthropological references. For estimates of times, those values for louse represents louse generations (g), and the values between [] for hominins are in years (y). For this table, louse values were estimated considering an infestation of 20 lice in 30% of hominin population and a generational time of 18 days.

| Details/Historic Events | Parameter | Distribution | Minimum-Maximum | References |
| --- | --- | --- | --- | --- |
| *Effective population* |  |  |  |  |
| Louse Ancestral Population<br>[Ancestral hominins] | N<sub>A</sub> | Uniform | 1,800-36,000<br>[300-6,000] |  |
| Louse cluster I<br>[AMH] | N<sub>1</sub> | Uniform | 1,800-36,000<br>[300-6,000] |  |
| Louse cluster II<br>[Neanderthals] | N<sub>2</sub> | Uniform | 1,800-36,000<br>[300-6,000] | [35] |
| Louse Hybrids<br>[Hybrids between AMH and Neanderthals] | N<sub>3</sub> | Uniform | 270-5,400<br>[?] | 15% hybrids from current study |
| *Time Events\** |  |  |  |  |
| Time of split clusters I and II<br>[Split between AMH and Neanderthals] | t<sub>s</sub> | Uniform | 1,200,000 - 16,000,000g<br>[600,000-800,000y] | [35] |
| T1 ancestral polymorphism from cII<br>[Presence of Neanderthals in Mongolia] | T1 | Uniform | 2,400,000 - 2,800,000g<br>[120,000 - 140,000y] | [36]<br>[38] |
| T2 ancestral polymorphism from cI<br>[Presence of AMH in Middle East] | T2 | Uniform | 1,000,000 - 1,600,000g<br>[50,000-80,000y] | [27] |
| Time of ancestral admixture<br>[AMH-Neanderthals in sympatry] | t<sub>aA</sub> | Uniform | 1,000,000 - 1,600,000g<br>[50,000-80,000y] | [20] |
| *Times of Recent Admixture* |  |  |  |  |
| Time of admixture E1<br>[European colonization] | T<sub>aE1</sub> | Uniform | 2,000 – 20,000g<br>[100 – 1,000y] |  |
| Time of admixture E2<br>[Migration after World Wars] | T<sub>aE2</sub> | Uniform | 800 – 2000g<br>[40 – 100y] |  |
| Time of admixture E1<br>[Initial Globalization] | T<sub>aG</sub> | Uniform | 200 – 800g<br>[10 – 40y] |  |
| Time of admixture E1<br>[Current time -t sampling] | T<sub>aC</sub> | Uniform | 20 – 200g<br>[1 – 10y] |  |

*Admixture rate*
|  |  |  |  |
| --- | --- | --- | --- |
| Rate of admixture | ar | Uniform | 0.001-0.999 |

*Mutation rate*
|  |  |  |  |
| --- | --- | --- | --- |
| Tetranucleotide microsatellite mutation rate | Um1 | Loguniform | $10^{-9}$–$10^{-3}$ |
| Trinucleotide microsatellite mutation rate | Um2 | Loguniform | $10^{-7}$–$10^{-3}$ |
| Dinucleotide microsatellite mutation rate | Um3 | Loguniform | $10^{-6}$–$10^{-2}$ |

\*Time parameters for lice are expressed as the number of generations of lice back in time assuming a generation time of 18 days [73].
The values for hominins refer to years.

For each model, we considered three levels of parasitism: a low level of parasitism with 10 lice per head and only 10% infested (S2 Table); an intermediate level, with 20 lice per head and only 30% infested (Table 2); and a high level of lice, with 50 lice per head and 50% infested (S2 Table) [71].

The louse life cycle includes egg, three nymphal stages, and adult. Therefore, each demographic model and each level of parasitism, was tested for three different generational times to account for natural length variation that could occur as was reported in the literature. For this study we considered the time of head lice maturation from egg to reproductive male adult and egg-laying-adult female of 18 days, 27 days and 36 days (Table 2, S3 Table) [72–74]. All microsatellites used the default generalized stepwise mutation model and were assigned a log uniform distribution with mean mutation rates between 10^-9^–10^-2^. The standard mutation rate for microsatellites is considered to be between 10^-6^–10^-2^ per locus per generation [75]. However, due to the possibility of different repeat lengths having different mutation rates [76] for microsatellites, tetra-, tri- and di- nucleotide repeats were modeled separately. Thus, preliminary runs were carried out in order to optimize mutation rates [77] following Bertorelle et al. [78]. A range in allele size of 40 contiguous allelic states was used, as recommended by the program developers (A. Estoup, personal communication).

Summary statistics calculated for microsatellite loci were as follows: mean number of alleles, mean genetic diversity, mean size variance, and mean *F*_ST_. The selection of summary statistics was empirical based on preliminary tests where these summary statistics were used and able to recover the models and parameters following Bertorelle et al. [78]. Through these preliminary tests the above summary statistics demonstrated their suitability for our data. We performed about 714,000 simulations using prior parameters for each scenario, thereby producing a reference table comprising 5,000,000 simulated data sets. First, we pre-evaluated the fit of observed values to prior distributions of scenarios, by conducting a Principal Components Analysis (PCA) using data sets simulated with the prior distributions of parameters and the observed data as well as data sets from the posterior predictive distribution to check the suitability of the model (scenarios and prior parameters). The posterior probability of each scenario was assessed by performing direct approach and a polychotomous weighted logistic regression that estimates the difference between simulated (from the reference table) and observed data sets [79,80] after linear discriminant analysis on summary statistics [70]. Only the posterior probabilities for the 1% (50,000) of simulated data sets closest to the observed data were retained. Confidence in choice of a scenario was estimated as the proportion of times the selected scenario did not possess the highest posterior probability compared to competing scenarios for the 500 simulated data sets (type I error). Confidence in scenario choice was further tested including the estimation of the prior error rate, which is calculated as the probability of choosing an incorrect model when drawing model index and parameter values into priors (the probability of selecting the chosen scenario when it is not correct, a type-II error). After scenario choice, we proceeded to parameter inference estimated from the modes and 95% confidence intervals (CI) of their posterior distributions.

## RESULTS

### Nuclear Genetic Diversity

The 274 human louse specimens comprising 25 geographic sites through 10 worldwide geographic regions revealed high microsatellite diversity with overall five to 20 alleles per locus and an average of 12 alleles per locus (Table 1, S1 File). In the 11 sites with more than 8 lice analyzed, H_O_ values ranged from 0.032 to 0.524 (average of 0.257) and H_E_ values ranged from 0.248 to 0.706 (average of 0.402), reflecting a general pattern of deficit of heterozygotes (Table 1). Furthermore, in all 11 louse populations, genotype proportions deviated significantly from Hardy-Weinberg expectations with values of *F*_IS_ significantly positive and an overall *F_IS_* value of 0.469 (*P* < 0.00001) (Table 1). Overall, there were high values of genetic differentiation among the 11 sites, with *F*_ST_ values of up to 0.663 between the sites: Mongolia (Mn) and Netherlands (Neth) (S4 Table). The lowest *F*_ST_ of 0.053 was found between sites in New York (Oce) and Tennessee (Nash).

### Mitochondrial diversity

We also obtained the COX1 sequence data from 175 head lice spanning 18 sampling locations throughout nine geographic regions (Table 3). One hundred four (104) lice belonged to haplogroup A and 71 to haplogroup B (Table 3). As expected based on previous studies [17,39,40], haplogroup A shows a worldwide distribution including Africa, Asia, Europe and the Americas. For our sampling, haplogroup B has a narrower distribution and is found only in lice from Europe, North and Central America (Table 3).

**Table 3.** Nuclear and mitochondrial louse genetic diversity. Geographic distribution of 175 human lice including combined information of their mitochondrial DNA haplogroups based the COX1 gene and their assignment to each louse major nuclear genetic clusters described in this study (GenBank Accession numbers: KF250510-KF250546 [17] and OR072805-OR072814 [current study])

| Geographic Region | Group | Country | Site | mtA+clusterI | mtB+clusterI | mtA+clusterII | mtB+clusterII |
| --- | --- | --- | --- | --- | --- | --- | --- |
| Africa | AIS | Rwanda | Rw | 1 |  |  |  |
| South Asia | SAs | Nepal | Np | 1 |  |  |  |
| Southeast Asia | SAs | Cambodia | Cam | 21 |  |  |  |
|  |  | Laos | Ls | 1 |  |  |  |
| East Asia | EAs | Mongolia | Mn | 6 |  |  |  |
| Europe | E | Turkey | Tur |  |  |  | 7 |
|  | E | Croatia | Croa |  |  |  | 5 |
|  | E | Norway | Nw |  |  | 1 |  |
|  | E | Netherlands | Neth |  |  | 49 | 5 |
|  | E | France | Fr |  |  | 2 |  |
|  | E | United Kingdom | UK |  |  |  | 26 |
| North America | NA | USA | Oce |  |  | 12 | 4 |
|  | NA | USA | SF |  |  | 1 |  |
|  | NA | USA | Nash |  |  |  | 3 |
|  | NA | USA | MC |  |  | 1 | 9 |
| Mesoamerica | MA | Mexico | MX |  |  | 6 |  |
| Central America | CA | Honduras | Hon |  | 12 |  |  |
| South America | SAm | Argentina | Arg |  |  | 2 |  |
| Total |  |  |  | 30 | 12 | 74 | 59 |

### Nuclear genetic clusters at the worldwide scale

All clustering methods: STRUCTURE, PCoA and DAPC were concordant in identifying the distinct nuclear genetic clusters: Cluster I (cI) with subclusters Ia and Ib and Cluster II (cII) with subclusters IIa and IIb (Fig 1; S1 Fig). The differences in their geographic distribution can be seen where cI defined by head lice from Africa, Israel, Asia and the Americas comprised a more worldwide distribution, while cII composed of head lice mainly from Europe and the Americas reflects a more restricted geographic distribution. While most of the genetic diversity of human louse populations from around the world seems to be highly structured geographically, approximately 12% of the lice we sampled (33 of 274 lice) appear to be hybrids (lice with *q*-values less than 0.8), with 75% (25 lice) of the hybrids found in the New World.

In the STRUCTURE analysis, after a signal for *K* = 2 (Δ*K* = 858.853, S2 Fig), increasing *K* values in the STRUCTURE plots revealed geographically localized structure. At *K* = 3, a new cluster formed within Europe from the original cII, composed of lice primarily from the Netherlands forming the subcluster cIIb (light blue color). When *K* was increased to 4 a population from Honduras and other parts of the Americas emerged from cluster I as the Ib subcluster, with the other samples remaining in subcluster Ia. This Honduras subcluster (Ib) may be indicative of Native American louse lineage.

The Principal Coordinate Analysis (PCoA) supported the two main nuclear genetic clusters and subclusters identified by STRUCTURE, with putative hybrid lice not assigned to either cluster, rather placing hybrids between them (Fig 3). Finally, the eigenvalues obtained in the DAPC analysis showed that the genetic structure was captured mainly by the first principal component representing cI (to the right) and cII (to the left) (S1 Fig).

**Fig 3.** Human louse worldwide nuclear genetic diversity. A) Genetic clusters inferred from STRUCTURE simulations at *K* = 2: Cluster I (cI, Orange) and Cluster II (cII, Blue), *K* = 3 cI and cIIa-cIIb, and *K* = 4 cIa-cIb and cIIa-cIIb. In the bar plot, each louse is represented by a single vertical line and the length of each color segment represents the proportion of membership (*Q*) to each cluster. At the top of the STRUCTURE bar plot, an additional bar was added using different colors for each continent and main geographic region where each louse sample was collected. At the bottom of the STRUCTURE bar plot, the codes correspond to geographic origin from the lice using abbreviations as defined in Table 1. B) Distributions of points in the first two dimensions resulting from Principal Coordinates Analysis (PCoA) conducted using pairwise genetic distance comparisons. Each site was color-coded to match STRUCTURE results (e.g., sites in Asia were color-coded orange, and sites in Europe were color-coded blue). These codes correspond to geographic origin and abbreviations as defined in Table 1. Map author: Maulucioni (https://commons.wikimedia.org/wiki/File:World_map_with_the_Americas_on_the_right.png). This file is licensed under the Creative Commons Attribution-Share Alike 4.0 International license. https://creativecommons.org/licenses/by-sa/4.0/legalcode

Additionally, we conducted further analysis with all the dataset excluding hybrids as identified through STRUCTURE *q*-values. Thus, when using the reduced dataset only non-admixed individuals with shared ancestry *q*-values larger than 0.8 as suggested by [56],the STRUCTURE analysis reflected similar results identifying cI and cII, with their corresponding subclusters at *K* = 4 (S3 Fig).

### Worldwide cluster I

All lice that belong to the cI (*q*-values larger than 0.5) were included in a second STRUCTURE simulation to assess the substructure within this cluster. At *K* = 2, this figure showed the presence of two clusters: one composed of lice from the Old World (Africa, Asia and Europe), and the other of lice from the New World (Americas) (S4 Fig). The further split of the genetic clusters during the increase of defined *K* in the STRUCTURE analysis showed at *K* = 3 the emergence of an Asian cluster except for most of the lice from Mongolia that remains within the Old Word cluster (S4 Fig). At *K* = 4, a subset of those lice from Mongolia formed a unique new cluster (S4 Fig).

### Geographically restricted cluster II

According to our current sampling, cII has a narrower geographic distribution than cI and is limited mostly to Europe and the Americas (Fig 3). This cluster showed a substructure at *K* = 2 where most of the lice from the Netherlands plus a few others form a distinct cluster (light blue in plot) (S5 Fig). Both these individual cluster analyses for cI and cII highlight the geographic structure observed among louse populations.

### Allelic richness analysis per main genetic cluster

We estimated the allelic richness per main non-admixed clusters I and II (without hybrids) using the rarefaction method implemented in the computer program ADZE (Allelic Diversity Analyzer) Version 1.0 [62]. Allelic richness per non-admixed cluster was higher in cluster I than II (S6 Fig). This pattern reflects very different evolutionary histories for the two clusters. While high allelic richness is consistent with a large effective population size, it could also be concordant with greater genetic structure. It is also noteworthy that clusters I and II differ in sample size (80 lice in the cI versus 174 in cII).

### Statistical analysis of genetic clusters and mitochondrial clades

The combined analysis of the nuclear cluster membership frequency and the mitochondrial haplogroup were plotted using a Correspondence Analysis (CA) and showed the presence of four well defined groups (S7 Fig). Dimensions 1 and 2 represented the largest amount of inertia or variance representing a measure of deviation of independence. Each one of the dimensions possesses an eigenvalue that represents its relative importance and how much of the variance is explained. In our study, one dimension represents 58.7 % of the inertia and two dimensions represents 99.6% of the inertia. Of this, dimension 1 differentiated both nuclear genetic clusters: cI and cII, while dimension 2 differentiated the mitochondrial haplogroups. One group corresponds to lice of nuclear cluster II-haplogroup A and includes lice from Europe, North America, Mesoamerica and South America. The second group is also nuclear cluster II-haplogroup B from the same regions, except South America. The third group is composed of individuals of cluster I-haplogroup A: South Asia, Africa-Israel-Spain, and some parts of Europe, while the last group belonged to cluster I with haplogroup B and was found in Central America (S7 Fig). Furthermore, multiple comparisons (non-parametric Kruskal–Wallis test) indicated that the distribution of the individuals assigned to the nuclear genetic clusters I and II were not distributed randomly among the eight geographic regions (H = 103.55, *P* < 0.0001). The cluster membership median values were significantly different among: South Asia (SAs), East Asia (EA), Africa-Israel-Spain (AIS), Central America (CA), and Europe (E) (*P* < 0.05). On the other hand, cluster membership medians values for North America (NA), South America (Sam), and Mesoamerica (MA) were not significantly different. Both nuclear genetic assignment and haplogroup frequencies differed significantly among the eight geographic regions (nuclear cluster assignment *Χ*^2^ = 175, *P* < 0.0001; haplogroup frequencies *Χ*^2^ = 42, *P* < 0.0001).

### AMOVA by nuclear clusters and mitochondrial clades

Hierarchical analyses of molecular variance (AMOVA) employed to explore the distribution of nuclear genetic variation considering the two nuclear genetic clusters: cI and cII revealed significant genetic differentiation between them (*Φ*_ST_ = 0.33; *P* < 0.00). When examining only the non-admixed individuals (those with at least 0.8 of coefficient of ancestry values) the differentiation between cI and cII increased (*Φ*_ST_ = 0.4; *P* < 0.00).

### Migration patterns considering nuclear genetic clusters and mitochondrial clades

The historical migration pattern was estimated in Migrate-n 3.3.0 using maximum likelihood inference and the Brownian motion mutation model [67,68]. Given that previous studies have suggested a different evolutionary history between the mitochondrial haplogroups A and B, we evaluated a full migration model with four populations defined by both the nuclear clusters inferred from STRUCTURE and the mitochondrial haplogroups: cI – mtDNA A; cI – mtDNA B; cII – mtDNA A and cII – mtDNA B. Migration rates between these four populations varied. In general, migration rates within nuclear clusters were high, contrasting with low migration rates within mitochondrial haplogroups (S8 Fig). For example, we observed the highest migration rates within cluster II (cluster II – mtDNA A and cluster II – mtDNA B) and some of the lowest migration rates between cluster I – mtDNA A and cluster II – mtDNA A. Because STRUCTURE analysis indicated the effect of geography on the genetic structure of human lice, further studies including more samples from each sampling site are needed to further understand migration patterns among geographic human louse populations.

### Demographic history

The presence of two louse distinct nuclear genetic clusters (I and II) could be the result of the retention of ancient polymorphisms due to incomplete lineage sorting (ILS) or admixture from previous isolated lineages. We tested seven different demographic models in which each louse genetic cluster and its hybrids evolve on different human host populations and diverge in tandem with them using the Bayesian computation (ABC) method as implemented in the DIY-ABC package [52]. The models reflect different hypotheses that followed human evolutionary events from the admixture between Neanderthals and AMH (Models 1-3) to human movement around the globe (Models 4-7) (Table 2, Fig 2). We tested these 7 demographic models using three different levels of louse parasitism as well as three different generational times creating a total of 63 scenarios (S9 Fig). The three most strongly supported demographic models (5, 6 and 7) were topologically similar with each involving admixture between the two nuclear clusters at different times since migrations after World Wars (Model 5), initial globalization (Model 6) and current times (Model 7) (Fig 4, Table 4, S9 Fig). Considering 10% and 30% levels of louse parasitism, for any of the generational times (18 days, 27 days or 36 days), the comparison of the seven demographic models showed Model 7 (current time) as the model with the highest logistic posterior probability of ca. 0.90 (Fig 4, Table 4, S9 Fig). For the high level of parasitism at 50%, the longer the generational time the older admixture models were shown with higher probabilities (S9 Fig). For example, when the longest generational time of 36 days was used in the scenario, then Model 4 appeared as the scenario with the highest probability. This model corresponds to the European colonization of the New World. When 27 days were chosen as generational time, Model 5, dated around the migrations after World Wars, presented the highest probability. Finally, when the generational time was set at 18 days, Model 6 that it is associated with Globalization shows the highest probability (S9 Fig).

**Fig 4.** Posterior probabilities for each of the seven demographic models. Graphs of posterior probabilities based on DIY-ABC analysis comparing the seven demographic models when considering intermediate louse infestation level where 30% of the hominin population have lice with 20 lice per head. A) The posterior probabilities are estimated using the direct approach and in B) they are inferred by a logistic approach [70]. Each scenario was color-coded as follows: Model 1 – green, Model 2 – red, Model 3 – light blue, Model 4 – pink, Model 5 – yellow, Model 6 – black, Model 7 – gray.

**Table 4.** Posterior probabilities for the seven demographic models shown in Fig 2. Probabilities obtained through the logarithmic method (with 95% confidence intervals in brackets) for the seven scenarios inferred from DIY-ABC analysis. For each model three parasitism levels were considered where 10% indicates that 10% of the hominin population have lice with 10 lice per head, 30% and 50% indicate that 30% and 50% of the hominin population have 20 and 50 lice per head, respectively. Generational time was set to 18 days. For each comparison, the selected scenario (bold entry in shaded cell) was the one with the highest probability value. Confidence in scenario choice, shown under probability values, was evaluated by estimating the type I error rate from 1000 randomly constructed simulated data sets, for Models 5 and 7, respectively.

| Models | Parasitism Level |  |  |
| --- | --- | --- | --- |
|  | 10% | 30% | 50% |
| 1: Retain Polymorphisms from cI | 0.0000 [0.0000,0.0000] | 0.0000 [0.0000,0.0715] | 0.0000 [0.0000,0.1130] |
| 2: Retain Polymorphisms from cII | 0.0000 [0.0000,0.0000] | 0.0000 [0.0000,0.0715] | 0.0000 [0.0000,0.1130] |
| 3: Ancestral admixture cI and cII | 0.0000 [0.0000,0.0000] | 0.0000 [0.0000,0.0715] | 0.0000 [0.0000,0.1130] |
| 4: European colonization of the New World | 0.0000 [0.0000,0.0000] | 0.0001 [0.0000,0.0716] | 0.0560 [0.0000,0.1522] |
| 5: European migration to the New World after WWs | 0.0000 [0.0000,0.0000] | 0.0071 [0.0000,0.0757] | **0.5204 [0.4477,0.5930]** |
| 6: Globalization | 0.0026 [0.0011,0.0041] | 0.3792 [0.2622,0.4963] | 0.2754 [0.2042,0.3467] |
| 7: Current time - sampling | **0.9974 [0.9959,0.9989]**<br>0.335 | **0.6136 [0.4959,0.7312]**<br>0.317 | 0.1482 [0.0095,0.2869]<br>0.372 |

## DISCUSSION

In this study, we found that the nuclear microsatellite data from lice were grouped in two distinct genetic clusters: cI and cII. Cluster I had a worldwide distribution whereas cII was restricted to Europe and the Americas (Fig 5). Remarkably, the two nuclear clusters revealed a significant genetic differentiation between them (*Φ*_ST_ = 0.33; *P* < 0.00) despite having geographic overlap in either Europe or the Americas. In the Structure analysis at *K* = 4 each of the two Clusters further split into subclusters: Ia-Ib and IIa-IIb, which were sorted based on their geographical origin (i.e., continents). These results are consistent with our previous work where we assessed louse microsatellite genetic diversity in fewer samples from only four geographic regions (North America, Central America, Asia, and Europe) [16]. Both this current study and our past work support the idea that head louse populations are strongly structured geographically. This current study highlights a genetic relationship between Asia (cIa) and Central America (cIb) subclusters supporting the idea that the Central America cluster (cIb) is probably of Native American origin since a potential source population for the first people of the Americas has been suggested to be East Asia [81–83] (Fig 5). Thus, microsatellite data suggest that current human louse populations in the Americas retain human louse genetic diversity from louse brought by the first people [16]. This work adds to our previous studies showing that human lice could be used to track aspects of human evolution.

**Fig 5.** Proposed global co-migration of human lice and humans. Top: Map depicting the collection sites of the human lice included in this study. The color of each circle corresponds to the majority nuclear genetic cluster to which sampled individuals were assigned. Sites with admixed lice are indicated with patterned circles including colors of the two major genetic clusters at that site. The proposed migrations of anatomically modern humans out of Africa into Europe, Asia and the Americas, as well as the more recent European colonization of the New World are indicated with thick grey arrows. Hypothetical human louse co-migrations are indicated with orange and blue arrows. At the bottom, the STRUCTURE plot from Fig 3A corresponding to the assignment of 274 lice from 25 geographical sites at *K* = 4 (Table 1) is shown. Map author: Maulucioni (https://commons.wikimedia.org/wiki/File:World_map_with_the_Americas_on_the_right.png). This file is licensed under the Creative Commons Attribution-Share Alike 4.0 International license. https://creativecommons.org/licenses/by-sa/4.0/legalcode

Other human parasites have also shed light into different aspects of human evolution [11–13]. For example, the genetic diversity of a bacterial parasite, *Helicobacter pylori*, which infects the stomach of humans, has been studied to provide insights into human migrations and strong correlations between the parasite and human host have been found reflecting the out of Africa expansion [84,85], the colonization of the Austronesian speaking region [86], and the settlement of the Americas by Asians across the Bering Strait [87]. However, some microorganisms such as bacteria and viruses that reproduce clonally could also be affected by selective sweeps that could lead to the removal of entire lineages, or to gene specific as well as genome wide reductions in genetic diversity. For those scenarios where parasites have clonal reproduction, the effect of selection could be strong resulting in parasite phylogenies reflecting the evolutionary history of their genes under selection (selective sweep) rather than the demographic history of the host. On the other hand, macroparasites (such as lice) often reproduce sexually, which lessens the effect of selective sweeps since it would only affect closely linked loci in their genomes [88]. Thus, sexually reproductive parasites may track host evolutionary history more accurately than clonal parasites. For this reason, the use of macroparasites, such as pinworms [89] and lice [2,3], that are also obligate parasites and co-evolved with their host, has become increasingly common in the study of human evolution. In fact, the human louse, *Pediculus humanus*, is one of the most well studied human parasites [1–6,14–19,21–28].

Human lice can disperse from a host to another at any motile stage (nymphs and adults), however adults are more likely to transfer from one head to another [90]. Lice can be transferred via direct body contact, for example during a fight, living with others, or through any other means where two heads are in close proximity to each other [91]. Additionally, head lice can be transferred via indirect contact, such as through sharing clothes that cover the head or combs. In this case the shared item may still have living lice that can be transferred to the non-infested person [92]. Louse ability to transfer from host to host directly or indirectly along with louse life history traits such as larger population sizes and faster rates of molecular evolution compared to their hosts reveal cryptic aspects of their hosts including how the host interact with other species, and host cryptic population structure. These host aspects might not be evident using the host data alone or not yet discernible in the host’s genome [9,93,94]. Differences between parasite and host intrinsic life-history traits may lead to incongruities in patterns of population structure between hosts and their parasites [95–99]. This phenomenon has been observed amongst several parasites including flies [99], helminths [100], and feather lice [101]. In one case, coevolutionary studies of the relationship between Arctic Galliformes and their parasitic lice in Alaska suggest lice consistently move among different host species and populations in that region [102]. While the bird hosts present some level of population structure and/or isolation by distance, the parasitic lice do not show any structure [102]. This lack of congruence between the genetic structures of the parasite and host suggests that there are still some cryptic contact or distributional overlaps among host bird populations. Although, these species interactions were not perceptible from the host data alone, they could be useful for ecological and conservation efforts. In our current study, we show the presence of two distinct nuclear genetic clusters: I and II, which are not observable in the human host. These could be the result of the retention of ancient polymorphisms due to incomplete lineage sorting (ILS) or admixture from previous isolated lineages. To test if current louse genetic diversity is the result of ILS or admixture we used the approximate Bayesian computation (ABC) method as implemented in the DIY-ABC package [70].

We performed ABC analyses for seven demographic models, three different levels of parasitism and three different generational times (Table 2, 4, Figs 2, 4, S9 Fig). The demographic model with higher probabilities under 10% and 30% parasitism level corresponded to the model suggesting recent admixture between the two nuclear Clusters I and II in Model 7. This model supports the STRUCTURE analysis, where most of the hybrids are being observed in the New World. Although for cases of higher parasitism, older admixture models were supported including models 4, 5, and 6. It could be possible that in the past both infestation rate and generational time could have been different due to different climate and human environmental conditions. In fact, several studies have shown that louse life history traits changed due to temperature. For example, it was reported that under laboratory conditions oviposition, egg hatchability, and fertility drastically varied with temperature suggesting that louse population sizes might increase in the winter months and decrease in summer [103]. Another study reported that *P. h. humanus* ceased to lay eggs at temperatures beneath 20°C and lay rapidly at 37°C. These conditions lead to a mean life span of 16 days and 27 days, respectively [72]. Despite the variation in the life cycle in relation to temperature, head lice were recorded from every group of man in every part of the world, even where it is known that lice are rare [104]. This could be generally attributed to human habits rather than to the climate [105]. For the remainder of this discussion, we will focus on the 30% level of parasitism’s scenarios as it most likely reflects current louse population conditions [106–108].

While most of the genetic diversity in human louse populations around the world seem to be highly geographically structured, there are a total of 33 hybrids out of 274 lice analyzed, reaching 12% of our current sampling. However, 25 of those hybrids (75%) are found in the New World, which could represent a zone of secondary contact as a consequence of human colonization. The three models that consider recent admixture events from European migration after both World Wars (Model 5), initial globalization (Model 6) and current times at sampling (Model 7) were the most supported (Table 2, 4, Fig 4, S9 Fig). Thus, a new and puzzling insight from this study was the finding of such low prevalence of hybrid lice and that these hybrids are geographically concentrated in the New World. This pattern could reflect human colonization of the New World by Europe as mentioned above. However, if these colonization events have been occurring for many louse generations, this does not explain the low occurrence of hybrids. It could be possible that our sampling is spatially/geographically biased as many samples came from the Americas (43%) (Table 1). In that case, it could be possible that hybrids are more common in other regions than the current data reveals. To answer this question, future studies on genetic diversity of human lice should consider sampling other geographic areas in more depth. However, the lack of hybrids in the Americas deserves some further discussion. While this is beyond the scope of the present paper, we hypothesize that this pattern of limited admixture could be indicating genetic isolation between cI and cII such that the hybrids are not able to persist for many generations within the human louse populations. Thus, it is worthwhile to enumerate the potential causes for future research.

One potential mechanism of reproductive isolation may involve the effect of endosymbiont on host biology. Endosymbionts could also alter host biology in diverse ways, including the induction of reproductive manipulations, such as feminization, parthenogenesis, male killing and sperm–egg incompatibility, which could lead to reproductive incompatibility [109]. *Wolbachia* is a common intracellular bacterial symbiont of arthropods and filarial nematodes that has been shown to infect human lice [110–112]. When populations of insect hosts that contain different strains of *Wolbachia* interbreed, or when one population has *Wolbachia* and the other population does not, the matings (“hybrids”) resulted in reduced fitness or the complete absence of progeny. If this process were to occur in human lice, it could explain the low number of hybrids; however, because *Wolbachia* endosymbiont is transmitted with the mitochondrial genome through the egg cytoplasm, endosymbionts and mtDNA should reflect a similar history. In our study, mitochondrial lineages were grouped in two mitochondrial haplogroups. We found individuals of both the A and B haplogroups in both nuclear clusters (cI and cII). This is consistent with our initial findings [16] in that there is no one-to-one correlation between mitochondrial haplogroups (A and B) and nuclear genetic clusters.

The human louse, unlike most animals, which have single-chromosome mitochondrial (mt) genomes [113], has fragmented mt genomes, each with 9-20 minichromosomes [114,115]. The human head louse (*Pediculus humanus capitis*, family Pediculidae) and human body louse (*Pediculus humanus corporis*) have the most fragmented mt genomes with 20 minichromosomes with a conserved common pattern whereas each of the other protein-coding and rRNA genes has its own minichromosome [114]. Furthermore, this pattern of gene distribution and the fragmented mt genome apparently evolved in the MRCA of the human lice and have been retained since [116]. Because stretches of identical sequences (26–127 bp long) have been found to be shared between genes on different minichromosomes, this was thought to be evidence for inter-minichromosome recombination [114] however the frequency of recombination has not been clearly established. Although, there could be some concern about mt minichromosome recombination since we only focused on one gene, the probability of that affecting our results is expected to be minimal.

Insect species with nutritionally incomplete diets (e.g., phloem or blood), such as human lice often harbor mutualistic bacteria that synthesize missing nutrients [117–119]. As *Wolbachia*, these endosymbionts are intracellular, and are transmitted vertically from mother to offspring. The human lice endosymbiont is *Candidatus* Riesia pediculicola [120], and based on genomic studies, this endosymbiont carries a small plasmid needed for vitamin B5 synthesis [121] suggesting that this bacterial symbiont supplies the lice with vitamin B-complex, which the louse cannot produce by itself [122,123]. Studies have confirmed that this obligate human louse endosymbiont is housed in the mycetome, which is localized on the ventral side of the louse midgut. In females, this endosymbiont also migrates to the lateral oviducts at the beginning of oogenesis, which is transovarially transmitted to the progeny confirming its maternal inheritance [124,125]. In Boyd et al. [121] they compared the genomic sequence of *Candidatus* Riesia pediculicola found in lice from Cambodia (cI-mtDNA A) and the Netherlands (cII-mtDNA A) with the lab strain, and they found no indels between these two genomes. However, when they included the genome obtained from louse from Honduras (cI-mtDNA B), the authors found indels [121], reflecting that correlation between mtDNA clades and the endosymbiont. A recent study using a larger number of lice has also shown the codivergence of the endosymbiont *Candidatus* Riesia pediculicola and the mt clades of their *P. humanus* host [124,125]. In our study, since there is no correlation between the mitochondrial DNA and the nuclear genetic clusters and since *Candidatus* Riesia pediculicola co-diverge with the mtDNA lineages, this endosymbiont does not explain the low prevalence of louse hybrids.

An alternative explanation could be related to an unusual paternal genome elimination (PGE) that has been reported in other members of Psocodea particularly Liposcelididae and Phthiraptera where for these two lineages PGE has been suggested as the mode of sex determination [126]. Earlier studies have shown that some, but not all, male of the clothing lice, *P. h. humanus*, only transmitted their maternal copy of microsatellite markers while females showed a Mendelian gene transmission [127]. This mode of non-Mendelian inheritance by male lice is consistent a type of pseudohaplodiploid reproduction known as paternal genome elimination (PGE), in which males only transmit maternally inherited alleles to their offspring [128]. A more recent study [129] assessed this phenomenon in both clothing and head lice. In both ecotypes of *P. humanus* louse males exclusively (or, in some cases, preferentially) transmit only the maternally inherited alleles to their offspring revealing a genome-wide male transmission ratio distortion for both ecotypes in agreement with early findings from McMeniman and Barker (2006) [127]. In our study, out of the 33 hybrids, 19 were female, 10 were male, and 4 were juvenile. Under the PGE scenario, all 10 male hybrids would only pass their maternally inherited alleles to the offspring thus lessening admixture in the populations. Since for our study lice were collected at a single time point, there is no data of louse pedigree which would have been needed to address PGE in detail. In addition, as human louse sex ratios are female-biased, PGE would have a lesser impact. Future studies including louse family pedigree and the analysis of allele transmission patterns are needed to understand if the low number of hybrid lice is a result of PGE.

If hybrids present some type of phenotypes that are not well adapted compared to the non-admixed parental lice, this could lead to an alternative hypothesis. For example, epigenetic mechanisms like DNA methylation and histone modification are well known to influence gene regulation, phenotypic plasticity, development, and the preservation of genome integrity. When most of the studies of epigenetics are related to intra-generational times, there is a growing number of studies focusing on transgenerational epigenetics: the inheritance of a modified phenotype from the parental generation without changes in genes or gene sequence. This transgenerational epigenetics suggest that there are direct and indirect contributions of epigenetics to evolutionary processes [130]. One of the modes for transgenerational epigenetic inheritance is through “genomic imprinting” for which a gene is distinguished as patrigene or matrigene, depending on its paternal or maternal inheritance [131]. This mode is often referred as parent-specific gene expression (PSGE) and has been well studied among social insects, where it is shown that epigenetic changes are functionally responsible for caste formation [132]. Among lice, epigenetics has not been studied and opens an exciting new area of research. There could also be a relationship between PGE and PSGE that could be interfering with the viability and/or fecundity of hybrid lice, a question that warrants epigenomic studies in human lice.

### Final remarks

In this work, we assessed the use of human louse nuclear genetic diversity as a tool to reconstruct human migration around the world, by analyzing microsatellite data from 274 lice. In addition to well-known genetic analysis methods, we developed a new approach to evaluate demographic scenarios through DIY-ABC models by using data from the human host. This novel approach could also guide the development of new analyses in other host-parasite systems. Our results showed the presence of two divergent nuclear louse genetic clusters, with very limited and recent admixture, mostly in the New World. We suggested that this pattern is reflective of human migration to the Americas, with an early wave of louse-human co-migration during the population of the New World followed by the most recent European migration. We also hypothesized that there is a potential mechanism that is preventing admixture between these two divergent nuclear clusters, most likely epigenetic in origin. Due to the use of microsatellites that are known to be fast-evolving markers, our analyses are more suited to recent events, and slower evolving markers could provide insights into more ancient events. Further studies in human lice including more samples, whole genomic, and epigenomic approaches could provide new knowledge about louse evolution as well as their human host.

## Supporting information

Answers.to.Reviewers.2February2023

## Acknowledgments

We would like to thank former University of Florida undergraduate students: Gebreyes Kassu, Jackie Fane, and Lauren Justice for their help in the lab and former UF postdoc Marie de Gracia Coquerel, currently at the Donald Danforth Plant Science Center for comments in early version of the manuscript. We greatly appreciate the contribution of Katie Shepherd (Founder & CEO of The Shepherd Institute for Lice Solutions), Marieta A. H. Braks (Laboratory for Zoonoses and Environmental, National Institute for Public Health and the Environment, Netherlands) and collectors worldwide for providing lice used in this study. We would like to thank Aida Miró-Herrans and Niyomi House (University of Florida, USA) and Jan Štefka (Institute of Parasitology, Biology Centre CAS, Czech Republic) for comments on an earlier version on this manuscript.

## Supporting information

**S1 Table. Detail of the human louse samples used in the current study.** Groups reflect their geographic distribution and STRUCTURE clustering: Africa-Israel-Spain (AIS), South and Southeast Asia (SAs), East Asia (EAs), Europe (E), North America (NA), Mesoamerica (MA), Central America (CA), and South America (SAm).

**S2_S3 Tables. Prior distributions of population sizes parameters and times of events for testing for the demographic scenarios described in Table 2 and Fig 2.** Between the brackets [] are the host values obtained from anthropological references as shown in Table 2 of the main manuscript. Louse values were estimated using anthropological references under different louse infestation levels as well as different generational times. **S2 Table.** Population size priors considering 10% and 50% louse infestation level. **S3 Table.** Louse time priors are expressed considering number of generations of lice back in time assuming a generation time of 27 and 36 days, respectively. The values for hominins refer to years.

**S4 Table. Genetic differentiation.** Louse pairwise genetic differentiation based on F_ST_ values from 15 microsatellite loci among sites where 8 or more lice were collected. Sample sizes are in parentheses following population name. Values below the diagonal indicate F_ST_ values, *p*-values are shown above the diagonal; * indicates significant at nominal alpha level of 0.05.

**S1 Fig. Discriminant analysis of principal components (DAPC).** Cluster membership of each louse is depicted by a different colour inside their 95% inertia ellipses. DA eigenvalues are shown in the inserted graphs at the bottom right, whereby the numbers of the discriminants plotted against each other are indicated by dark grey and the remaining discriminants retained for the analysis in light grey. A total of 10 axes were retained in this DAPC, this is referred to the n-dim function in the adegenet package, and indicates the number of retained DAPC axes, which is affected by both the number of PCA axes and DA axes retained.

**S2 Fig. Number of clusters.** ΔK statistic of Evanno et al. [39] from STRUCTURE analysis from Fig. 3. K = 2 (ΔK = 858.853).

**S3 Fig. STRUCTURE plot including only non-admixed lice (no hybrids) from both clusters I and II at K = 4.** An ancestry threshold value of *q*-values as suggested in [38] was used because it is efficient and accurate at differentiating between non-interbreed and hybrids. Only individuals with *q*-value between 0 and < 0.20 or > 0.8 and 1 are classified as non-admixed and were included in this analysis.

**S4 Fig. Cluster I substructure.** STRUCTURE plot including only non-admixed lice (no hybrids).

**S5 Fig. Cluster II substructure.** STRUCTURE plot including only non-admixed lice (no hybrids).

**S6 Fig. Allelic richness.** Allelic richness per nuclear genetic cluster (cI, orange and cII, blue) using the rarefaction method implemented in the computer program ADZE (Allelic Diversity Analyzer) Version 1.0 [44]. Because allelic richness measurements considered sample size, the rarefaction method allows the estimation of allelic richness for different random subsamples of size “g” from the populations [45,46].

**S7 Fig. Correspondence analysis (CA).** CA of the nuclear cluster membership frequency and the mitochondrial haplogroup of head lice.

**S8 Fig. Gene flow.** Full migration model with four units defined by both the nuclear clusters inferred from STRUCTURE and the mitochondrial haplogroups: unit 1) cI – mtDNA A; unit 2) cII – mtDNA A, unit 3) cI – mtDNA B; and unit 4) and cII – mtDNA B. Values above and below the arrows indicate migration rates, arrows indicated directionality.

**S9 Fig. Graphs of posterior probabilities for each of the 63 scenarios per level of parasitism and generational time.** Louse infestation included three levels: 10%, 30% and 50%, where 10% indicates that 10% of the hominin population have 10 lice per head, the 30% level considers that each head would have 20 lice and only 30% of the population is infested, while the 50% reflects high level of parasitism with 50 lice per head in 50% of the population. For this study we considered the time of head lice maturation from egg to reproductive male adult and egg-laying-adult female of 18 days, 27 days and 36 days. Each demographic model was color-coded as follow Scenario 1 – green, Scenario 2 – red, Scenario 3 – light blue, Scenario 4 – pink, Scenario 5 – yellow, Scenario 6 – black, Scenario 7 – grey.

**Supporting information document** including a) Protocol for collecting human lice developed by the Reed lab; b) Questionnaire on inclusivity in global research; and c) Microsatellite data

