## Supplementary material for "Nuclear genetic diversity of head lice sheds light on human dispersal around the world": Answers.to.Reviewers.2February2023

USDA Agricultural Research Service

February 2nd, 2023

Dear Editor,

Our manuscript titled: “Nuclear genetic diversity of head lice sheds light on human dispersal around the world” by Marina S. Ascunce, Ariel C. Toloza, Angelica Gonzalez-Oliver, and David L. Reed was submitted to be considered for publication in *PLOS ONE* as a research article and we had received the reviewers and your comments.

We are very grateful for the reviewers’ valuable and thoughtful comments. We have now addressed the reviewer’s concerns and the editorial comments as follows:

**Reviewer 1**

1) …Louse 18 day life cycle… At a minimum, please include a discussion justifying the choice of 18 days, or some discussion of the results sensitivity to this factor.
*Answer:* The reviewer brought up an interesting question that we didn’t consider in our initial version of the manuscript. We followed the suggestion and included this topic in the discussion in the revised version of the manuscript. In addition, we performed a new series of analysis to include longer generational times of 27 and 36 days as previous authors have reported length variations. [Material and Methods, Results, Discussion]

2) Microbial coinfection (lines 604-606). I don't quite follow why you arrive at the conclusion that Reisia cannot be selecting against hybrids. Is this because of Reisia is assumed to have maternal inheritance (fair assumption) like the mt genes (less fair assumption) and as the mt doesn't seem to explain lack of hybrids, neither would Reisia? Might be quite off base here but this section seemed to be missing a connecting thought in your argument so please explain.
*Answer: Candidatus* Riesia pediculicola is an obligate human louse endosymbiont that is transovarialy transmitted to progeny and housed in the mycetome, which is localized on the ventral side of the louse midgut. Migration is initiated by a stimulating factor associated with the adult moult. In females, the symbionts migrate to the lateral oviducts at the beginning of oogenesis, and in male adults, the stomach disc will degenerate over time [1,2]. These studies confirm that this endosymbiont has a maternal inheritance. A recent study has also shown the codivergence of the endosymbiont *Candidatus* Riesia pediculicola and the mt clades of their *P. humanus* host [3]. In our study, since we didn’t find a correlation between mtDNA clades and nuclear genetic clusters, it is plausible to assume the same for *Candidatus* Riesia pediculicola, thus this endosymbiont cannot explain the lack of hybrids. We clarified this in our current version of the manuscript.

3) Mt inheritance. Given that human lice (all Anoplura in truth) have multichromosomal mt genomes, there is a strong potential for independent inheritance of different mt genes as they are no longer linked in a single chromosome as in most animals. It is probably worth commenting on how this violation of 'typical' mt inheritance models

would affect your methods (I am thinking not significantly as it is just 1 mt gene analysed) and independent association of mt and nuclear lineages.
*Answer:* The topics of lice fragmented mitochondrial (mt) genomes, mt minichromosomes and the potential of mt recombination were included in the revised version of this manuscript.

4) PGE & methods. In a similar vein please enumerate which of the pop gen methods used here have diplo-diploid mendelian inheritance as background assumptions for that method? The reliability of the conclusions requires us to be sure that they aren't an artefact of inappropriate analyses, and given that PGE is such a significant part of louse inheritance it needs thorough discussion. I was very pleased to see this section in the discussion (lines 606-622) but I would prefer to see a thorough dissection of the issue.
*Answer:* We have not analyzed PGE per se as the type of sample done only collected lice at a single time point, there is no data of louse family pedigree and allele transmission patterns which would have been needed to address PGE. We included in the discussion as another hypothesis to test in future studies as a potential mechanism of lack of hybrids. This clarification was added to the discussion section.

5) The text has many instances of small typos or expression that makes it harder to read and so need thorough revision before final publication. E.g. line 42: "show expression over the last", line 54: "allow us to harness", line 330 "pattern", line 569 "reflect". Just a small selection but the paper deserves a fine revision for readability.
*Answer:* We have done a proof-reading of the manuscript.

6) Fig 2 is not readable in the reviewer copy and the differences between the scenarios is not obvious. I think that this is largely due to low resolution which may be on the journal's side of the review process rather than the submitted figure but please check on this one. It is rather central to the paper.
*Answer:* We re-did the figure to improve resolution.

**Reviewer 2**

1. In the Introduction, please provide more details about how this study is different from your previous study. It becomes clear later that the biggest difference is breadth of sampling (additional continents, etc.), but I think it would be best to include this information up front.
*Answer:* We have clarified this as suggested by the reviewer.

2. Consider adding more discussion about sampling bias. Is it possible how you obtain samples could dictate your findings? For example, are certain demographics more likely to provide samples of lice? Some additional information in the Methods could help too. I realize there are ethical considerations with sharing sample data from humans, but I think it is important to address this possibility as much as you can.
*Answer:* We have enlarged this topic in the discussion in regard to geographic bias. However, we don’t have data about demographics, there were not collection of personal data. In general, pediculosis is known to affect children more than adults, however that it is part of the parasitism dynamics. On the other hand, studies have shown that there

is no significant link between pediculosis and socioeconomic status [4,5]. But, head louse infestation prevalence varies according to time, space, cultural practices and sanitation [6–8].

(3) Consider running a ML analysis with bootstraps on the COX1 data.
*Answer:* We have followed Light et al. (2008) [9], where the authors suggested a distance-based neighbor-joining (NJ) analysis to verify clade membership based on phylogenetic groupings considering the strong differences between the clades A and B. We included this reference in the revised version of the manuscript, also to address the reviewer, we conducted a ML analysis, and we obtained a similar tree pattern as the NJ tree.

4. Please be sure to clarify when you are using microsats vs COX1 data. For example, it is unclear in the section beginning Line 180.
*Answer:* We have clarified this in the revised version of the manuscript.

5. Please provide more details regarding PCoA and DAPC. How many components did you retain for the DAPC? In addition, please justify why you used these approaches (vs. PCA, etc.).
*Answer:* Principal Coordinate Analysis (PCoA) is a multivariate technique that allows describing and plotting the major patterns within a multivariate data set (e.g., multiple loci and multiple samples). PCoA is a scaling or ordination method that starts with a matrix of similarities or dissimilarities between a set of individuals and aims to produce a low-dimensional graphical plot of the data in such a way that distances between points in the plot are close to original dissimilarities. Each successive axis of the plot explains proportionately less of the total variation, such that when there are distinct groups, the first 2 or 3 axes will typically reveal most of the separation among them. PCoA aims to summarize the overall variability among individuals, which includes both the divergence between groups (i.e., structured genetic variability), and the variation occurring within groups. On the other hand, DAPC is a clustering method that maximizes the between-group variance while minimizing the within-groups variance. DAPC results are presented as multidimensional scaling plots. Because it summarizes the genetic differentiation between groups, DAPC achieves the best discrimination of individuals into pre-defined groups. Thus PCoA and DAPC are complementary analysis. A total of 10 axes were retained in our DAPC analysis, that information was included in our current version of the manuscript.

PCA (Principal Component Analysis) and PCoA are similar and they both create components – also called axes or dimensions – that summarize the original information of the data in a reduced space. However, Principal Component Analysis (PCA) is a classical ordination method for raw data, while Principal Coordinate Analysis (PCoA) applies to a distance matrix. Thus, the starting point matrix of similarities or dissimilarities for PCoA is different from that of PCA, which starts with the initial data matrix (e.g., presence versus absence of alleles in molecular marker data).

6. Differentiating between ILS and ancestral admixture is a fairly big deal, so I would mention this in the Methods when you introduce the DIY-ABC program. You mention this in the Discussion, but I think it’s a major plus of using this program.

*Answer:* Thank you, we have included this part in the introduction and the methods.

7. How did you choose the levels of parasitism (e.g., Line 288)? It's difficult to evaluate whether these are reasonable numbers of "low," "intermediate," and "high." Can you provide some references that support your decisions? Perhaps reference #53 is sufficient, but this is unclear from the text.
*Answer:* We defined the levels of parasitism based on the data reported by several authors. In all the works, head lice were more common at younger age and were independently of social economic settings. For example, the foundational works of Nuttall 1917 [10] and Buxton 1946 [11] reported that infestations levels varied considerably and can be included into three categories (low, intermediate and high). Similarly, the worldwide review of the epidemiology of head lice published by Falagas et al. 2008 [5] showed the same pattern. In another words, human head could harbor few (1-10), medium (11-50) and high (>50) adult lice per head. Of this, a negative correlation between the number of lice per head and the infested human heads was found. It means that the heads containing the highest number of lice are rare while most of the cases are in the low to medium category. More recently, Toloza et al. [12] studied the epidemiology of head lice in elementary schools together with the mathematical modeling of the evolution of a head lice colony by using collected data and simulations. Mathematical model allowed authors not only to predict the growth a colony for different number of founders, but also to understand the influence of the timing of their transfer to the colony. The study showed a high statistically significance of the number of lice per head found in a real context and the number of human heads containing them. It also detected the pattern of the infestation categories mentioned above and further inspection of the data showed that in each of the classes where the clusters or categories were found, there was a severely infested child that could have acted as the source of the infestations observed, playing the role of a superspreader. In addition, similar infestation levels of pediculosis was a recurrent health problem among pre-Columbian populations as it is worldwide today [13]. Overall, the population frequency of head lice varies according to time, space, cultural practices and sanitation [6–8].

8. What about the hybrid lice that are not from the New World? Is there anything unique about those samples that indicates why there is hybridization? Or perhaps this is evidence that hybridization would be more widespread with a larger sample size outside of the New World?
*Answer:* Yes, we agree with the reviewer, enlarging the sampling in other geographic regions beside the New World could show a different pattern and hybrids could be frequent in other regions. We clarified this in our revised version of the manuscript to address this limitation and the potential of geographic bias of our sampling. [Discussion]

9. I think it's too strong to say that it is "probably epigenetics" that is preventing admixture (line 650). I would tone down this language.
*Answer:* We re-wrote this idea.

10. There are a few typos incorrect word usage. I suggest giving the manuscript another read-through to tighten up the writing. I’ve included some examples I noticed below, in addition to some other minor comments:
- Line 41: Perhaps “ideal system for coevolutionary studies”?
- Line 42: Please clarify what you mean by “expand.” Do you mean “span?”
- Line 45: It is unclear how archaeological findings of lice are tied to coevolutionary studies. Please expand.
- Line 47: Close host-parasite relationships don’t always result in reciprocal adaptations, and it can be difficult to demonstrate this in natural systems. I would say something like “could lead to reciprocal adaptations. It’s also important to note that coevolution and cospeciation do not always go hand-in-hand.
- Line 51: Consider a word other than “inextricably.” Perhaps “close”?
- Line 53: “relatively faster”
- Line 55: “allowed researchers to harness the parasites’”
- Line 74: Do you mean “hominins”?
- Line 94: Remove parentheses.
- Line 102: Define parameters and models for what?
- Line 134: Please specify the extraction modifications.
- Line 330: “pattern”
- Line 353: “comprise”
- Line 524: “lessens”
*Answer:* We have expanded, clarified and resolved the points and typos requested in the revised version of the manuscript.

- The font in Figure 2 is a bit small. Please make it larger.
*Answer*: We re-did Figure 2.

- Line 184: Why did you use 8 lice as the cutoff for FIS, HO, and HE?
*Answer:* This was a rule of thumb criterium as the less individual in a population could lead to higher degrees of error in those estimates. When for population studies using microsatellites researchers aim to reach at least 25-30 individuals [14], this was not possible for most of the louse populations included in the study due the human dependent situation. The potential sampling bias was expanded in the revised version of the manuscript.

- Figure 4: Please explain the difference between 4A and 4B in the caption.
*Answer:* In 4A) The posterior probabilities are estimated using the direct approach and in 4B) they are inferred by a logistic approach. This was added to the legend of Figure 4.

- What is the top bar in Figures 3 and 5? Does this represent human genetic structure, or just delimiting geographic structure?
*Answer:* The bar is indicating continents and geographic regions to help with locations of the louse samples. The legends of the figures were edited to clarify this.

**Editorial comments / Journal Requirements:**
1. Please ensure that your manuscript meets PLOS ONE's style requirements

*Answer:* We revised the manuscript to ensure it follows PLOS ONE's style requirements.

2. Upon re-submitting your revised manuscript, please upload your study's minimal underlying data set as either Supporting Information files or to a stable, public repository and include the relevant URLs, DOIs, or accession numbers within your revised cover letter.
*Answer:* We have uploaded the study dataset as Supporting information file.

3. Figures 1, 5 and Striking image in your submission contain [map/satellite] images which may be copyrighted … We require you to either (1) present written permission from the copyright holder to publish these figures specifically under the CC BY 4.0 license, or (2) remove the figures from your submission.

*Answer:* For Figure 1 and Figure 5 and striking image, we re-did the figures using a map that complies with the CC BY 4.0 license, detailed information has been added to the legend of the map.

4. Please review your reference list to ensure that it is complete and correct.
*Answer:* We have conducted a revision of the reference list.

5. [Request from editorial office received on 2 February 2023] Please state what role the funders took in the study. If the funders had no role, please state: "The funders had no role in study design, data collection and analysis, decision to publish, or preparation of the manuscript."
Please include this amended Role of Funder statement in your cover letter; we will change the online submission form on your behalf.
*Answer*: "The funders had no role in study design, data collection and analysis, decision to publish, or preparation of the manuscript."

Thank you for considering our manuscript for publication in *PLOS ONE*.

Sincerely,

Marina Ascunce

14. Hale ML, Burg TM, Steeves TE. Sampling for Microsatellite-Based Population Genetic Studies: 25 to 30 Individuals per Population Is Enough to Accurately

Estimate Allele Frequencies. Poon AFY, editor. PLoS ONE. 2012;7: e45170. doi:10.1371/journal.pone.0045170
