## Supplemental Tables for "Nuclear genetic diversity of head lice sheds light on human dispersal around the world"

**S2 and S3 Tables. Prior distributions of population sizes parameters and times of events for testing the demographic models described in Table 2 and Fig 2.** Between the brackets [] are the host values obtained from anthropological references as shown in Table 2 of the main manuscript. Louse values were estimated using anthropological references under different louse infestation levels as well as different generational times.

**S2 Table. Population size priors considering 10% and 50% louse infestation level.**

| Historic Events | | 10% Parasitism | 50% Parasitism |
|---|---|---|---|
| *Effective population* | N | Minimum-Maximum | Minimum-Maximum |
| Louse Ancestral Population | $N_A$ | 300-6,000 | 7,500-150,000 |
| [Ancestral hominins] | | [300-6,000] | |
| Louse cluster I | $N_1$ | 300-6,000 | 7,500-150,000 |
| [AMH] | | [300-6,000] | |
| Louse cluster II | $N_2$ | 300-6,000 | 7,500-150,000 |
| [Neanderthals] | | [300-6,000] | |
| Louse Hybrids | $N_3$ | 45-900 | 1,125-22,500 |
| [Hybrids between AMH and Neanderthals] | | [?] | |

**S3 Table. Louse time priors are expressed considering number of generations of lice back in time assuming a generation time of 27 and 36 days, respectively. The values for hominins refer to years.**

| Historic Events | | 27 days | 36 days |
|---|---|---|---|
| *Time of Events* | Time | Minimum-Maximum | Minimum-Maximum |
| Time of split clusters I and II | $t_S$ | 7,800,000 - 10,400,000 | 6,000,000 - 8,000,000 |
| [Split between AMH and Neanderthals] | | [600,000-800,000y] | |
| T1 ancestral polymorphism from cII | T1 | 1,560,000 - 1,820,000 | 1,200,000 - 1,400,000 |
| [Split between AMH and Neanderthals | | [120,000 - 140,000y] | |
| T2 ancestral polymorphism from cI | T2 | 650,000 - 1,040,000 | 500,000 - 800,000 |
| [Presence of AMH in Middle East] | | [50,000-80,000y] | |
| Time of ancestral admixture | $t_{aA}$ | 650,000 - 1,040,000 | 500,000 - 800,000 |
| [AMH-Neanderthals in sympatry] | | [50,000-80,000y] | |
| *Times of Recent Admixture* | | | |
| Time of admixture E1 | $T_{aE1}$ | 1,300 - 13,000 | 1,000 - 10,000 |
| [European colonization | | [100 – 1,000y] | |
| Time of admixture E2 | $T_{aE2}$ | 520 - 1,300 | 400 - 1000 |
| [Migration after World Wars] | | [40 – 100y] | |
| Time of admixture E1 | $T_{aG}$ | 130 - 520 | 100 - 400 |
| [Initial Globalization] | | [10 – 40y] | |
| Time of admixture E1 | $T_{aC}$ | 13 - 130 | 10 - 100 |
| [Current time -t sampling] | | [1 – 10y] | |
